# Distribution and structural diversity of Type IV internal ribosome entry sites

**DOI:** 10.1101/2025.05.12.653553

**Authors:** Katherine E. Segar, Madeline E. Sherlock, Jeffrey S. Kieft

## Abstract

Internal Ribosome Entry Sites (IRESs) are RNAs that facilitate cap- and end-independent translation initiation in eukaryotes. Type IV IRESs, which include the hepatitis C virus IRES, directly bind the 40S subunit and require only a subset of the canonical initiation factors to function. As the full diversity and species distribution of these IRESs was unknown, we sought to identify and classify their full architectural variation. Using a secondary structure homology-based search method, we identified 163 putative Type IV IRESs from viruses with diverse hosts and phylogeny, including the first example in a double stranded viral genome. Clustering analysis based on the presence and overall size of secondary structure elements yielded three distinct groups, differentiated by secondary structure expansions and deletions. Chemical probing of representative IRESs from each cluster validated the predicted secondary structures and in vitro translation assays showed that structural differences correlate with functional variation. Our findings reveal distinct structural adaptations and patterns within the Type IV IRESs that may influence IRES function and mechanism.

## INTRODUCTION

Eukaryotic translation initiation is a multi-step process dependent on a modified nucleotide cap on the 5’ end of the messenger RNA (mRNA) template and many eukaryotic initiation factor (eIF) proteins (Brito Querido et al. 2024). Internal Ribosome Entry sites (IRESs) are RNA elements that facilitate translation initiation independently of this cap or the 5’ end of the RNA, often using a subset or none of the eIFs (Martinez-Salas et al. 2018). An advantage of this streamlined mechanism is that when canonical translation initiation is repressed, some IRESs still function, enabling the virus to avoid global translational repression during a cellular immune response. Many types of IRES RNAs have been identified, with mechanisms and factor requirements that depend on their secondary and tertiary structure (Martinez-Salas et al. 2018; Abaeva et al. 2024).

Type IV IRESs (also known as class III) include the hepatitis C virus (HCV) IRES, which is the founding member and the most studied (Li et al. 2024; Mailliot and Martin 2018). The HCV IRES structure contains stem-loops, four-helix junctions, and a pseudoknot organized into two main secondary structure domains (domains II and III) (Lukavsky 2009). Each domain contains subdomains (IIa, IIb, IIIa-IIIf) (**Fig. 1A**), with the pseudoknot interaction between IIIf and downstream RNA and a long-range base pair in IIIe (Lukavsky 2009; Li et al. 2024). Magnesium-dependent global folding of the IRES RNA involves a local compaction of the junction regions, which helps spatially position the emerging stem-loops to make specific interactions with the translation machinery (Kieft et al. 1999).

**Figure 1.**
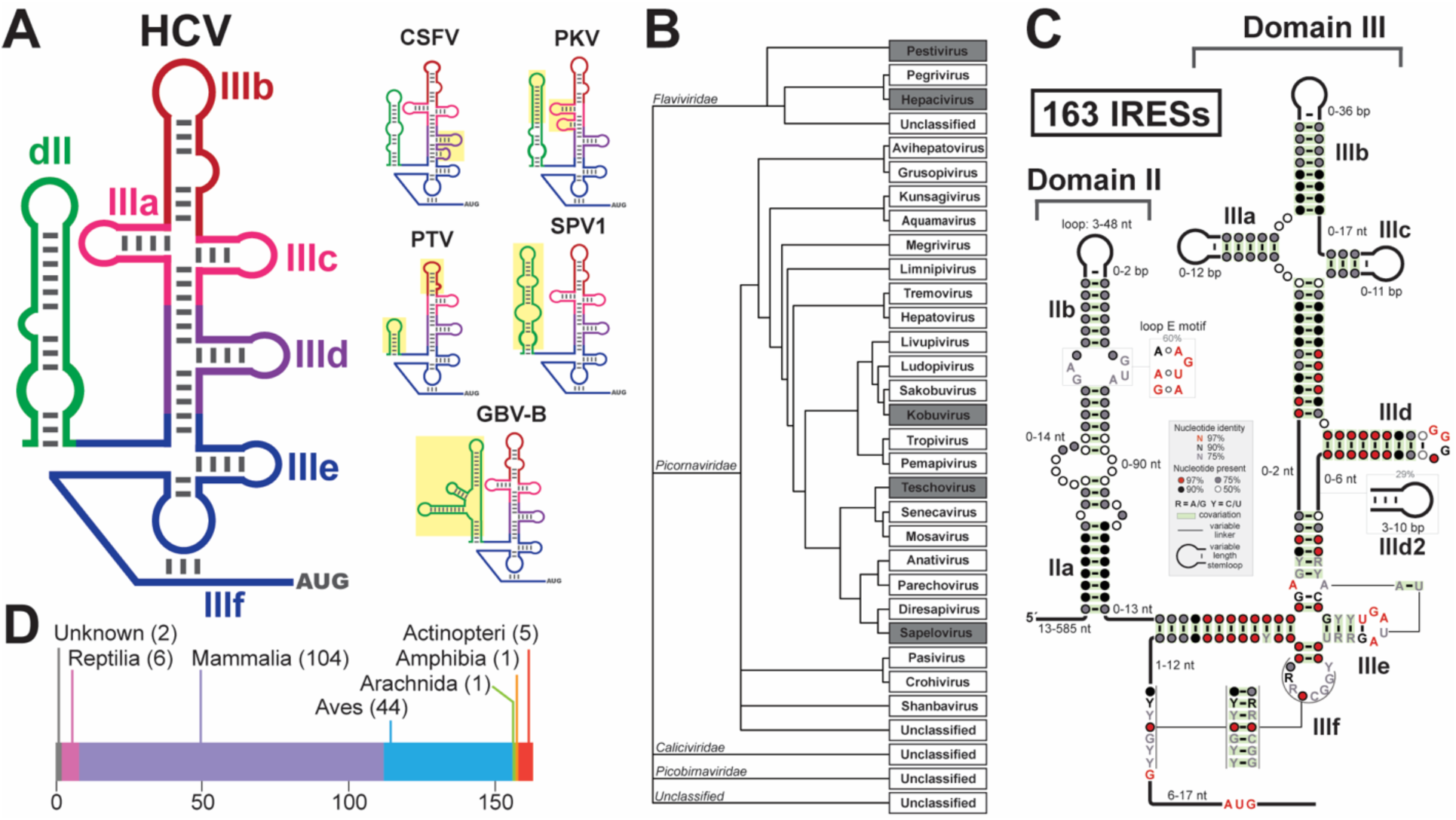
Identification of type IV IRESs. (**A**) Left: Cartoon secondary structure of the HCV IRES with domains labeled. Right: Cartoons of other Type IV IRES used in the seed alignment. Deviations from the HCV IRES structure are boxed in yellow. (**B**) Phylogenic tree of viral species with a Type IV IRES identified in the search. Grey boxes indicate species used in the seed alignment. (**C**) Consensus model using 163 curated sequences (see Table S1). Model generated using R-scape. (**D**) Distribution of the host organism of identified viruses.

The HCV-IRES does not require the cap-recognizing eIFs, rather the ribosome is recruited directly to the start codon through interactions between the IRES and the small (40S) ribosomal subunit (Pestova et al. 1998; Kieft et al. 2001). Key interactions are between the apical loop of IIId that base pairs with rRNA expansion segment 7 (Jubin et al. 2000; Kieft et al. 2001; Quade et al. 2015), and IIIa and IIIc that contact ribosomal protein eS27 (Hashem et al. 2013b; Quade et al. 2015). The HCV IRES also uses IIIb and IIIa to bind the multi-protein eIF3 complex (Sun et al. 2013; Sizova et al. 1998; Hashem et al. 2013b). During canonical translation initiation, eIF3 also directly interacts with eS27 (Brito Querido et al. 2020; des Georges et al. 2015), a site that is inaccessible when the IRES is bound to the 40S subunit. Structural studies of the HCV IRES and the related classical swine fever virus (CSFV) revealed that eIF3 is displaced from its normal binding site on the 40S subunit during IRES-driven initiation (des Georges et al. 2015; Hashem et al. 2013a; Sun et al. 2013). This suggests that the HCV IRES hijacks and remodels recycling 40S subunits – which contains eIF3, eIF1 and eIF1A – to access eS27, yet retains eIF3 (Sun et al. 2013; Hashem et al. 2013b; Jaafar et al. 2016). This complex then continues to form a preinitiation complex with eIF2 or eIF5B delivering initiator tRNA (Jaafar et al. 2016). The HCV IRES can begin its initiation process by binding each component of its complex sequentially or by binding an actively translating 40S subunit (Brown et al. 2022; Yokoyama et al. 2019), likely using some combination of these *in vivo*.

The HCV IRES is the prototype Type IV IRES, but others have been identified through sequence homology searches or by examining viral 5’ untranslated regions (UTRs) within phylogenic orders already known to contain them (Li et al. 2024; Asnani et al. 2015; Arhab et al. 2022). These strategies may overlook more structurally divergent IRESs or those residing outside expected viral clades. Identifying and testing more Type IV IRESs will lead to a much greater understanding of how IRES RNAs evolve and could create new tools and opportunities for biotechnology development. For example, a selection of Type IV IRESs could be used in therapeutic circular RNAs to drive translation at a desired rate in a specific context. This motivates an alternative strategy to identify and characterize more Type IV IRESs, potentially expanding the IRES toolkit for synthetic biology and RNA-based therapeutics.

To explore the full diversity of type IV IRESs, we utilized a proven bioinformatic approach with alignments of IRESs based on conserved secondary structural features and persistent sequence motifs. We identified 163 unique IRES sequences, including a viral clade not previously known to have IRESs. This expanded set was computationally clustered into three architecturally different groups that show a range of functional efficiencies; this likely reflects differences in how they interact with the translation machinery. This study expands the known examples of Type IV IRESs and catalogs their structural and functional heterogeneity.

## RESULTS AND DISCUSSION

### Homology-based search reveals new Type IV IRESs and phylogenic distribution

To fully catalog the structural diversity and phylogenetic distribution of Type IV IRESs, we used a homology-based bioinformatic approach that considers both sequence conservation and secondary structural patterns. We aligned six Type IV IRES sequences with biochemically verified secondary structures to create a ‘seed’ alignment (**Fig. 1A**). These six were the hepatitis C virus (HCV), classical swine fever virus (CSFV), porcine kobuvirus (PKV), GB virus B (GBV-B), simian picornavirus 1 (SPV1), and porcine teschovirus (PTV) IRESs (Brown et al. 1992; Fletcher and Jackson 2002; Fan et al. 2013; Rijnbrand et al. 2000; De Breyne et al. 2008; Chard et al. 2006). We used this seed alignment with the program Infernal to search a curated NCBI viral genomic database for RNA sequences that match the patterns found in the seed alignment (Nawrocki and Eddy 2013; Sayers et al. 2024).

The search identified 163 Type IV IRESs from a variety of viral species. While many were in previously identified viruses, a new putative Type IV IRESs was revealed in the *Picobirnaviridae* family (**Fig. 1B**). The *Picobirnaviridae* family includes double stranded RNA viruses and is a largely understudied viral clade (Delmas et al. 2019). This is the first reported Type IV IRES identified in a double stranded RNA virus. We also identified likely Type IV IRESs in currently unclassified viruses. All putative IRESs are found in the 5′ UTR of viral genomes, except for one that is found in the 3′ UTR. All identified IRESs are directly upstream of a subsequent open reading frame.

Previous comparisons of Type IV IRES structure were largely limited to a handful of examples at a time (Asnani et al. 2015). Our alignment created the opportunity to do a global assessment of secondary structural diversity and patterns to better understand the full architectural landscape of Type IV IRESs. Using this curated alignment, we created a covariance model that includes all putative Type IV IRESs, illustrating patterns of primary sequence and secondary structure conservation and variation (**Fig. 1C**) (Rivas 2020).

IIIdef is structurally conserved, consistent with it being the shared functional ‘core’ of the Type IV IRESs (Quade et al. 2015).. This includes the highly conserved ‘GGG’ in the apical loop of IIId that base pairs directly with (expansion segment 7) ES7 ‘CCC’ motif in rRNA (**Fig. S1A**) (Quade et al. 2015; Jubin et al. 2000). We found only 3 examples that had a ‘GGA’ or ‘GGC’ in place the canonical ‘GGG’ sequence (MN602325.1, MG600108.1, MK204421.1; Arhab et al. 2020), but this variation is not host species specific and the implications are unclear. The IIIf pseudoknot is also maintained although its sequence is variable, consistent with its role in positioning the start codon through non-sequence-specific interactions (Quade et al. 2015). Finally, a long-range base pair involving IIIe is conserved, consistent with its role in enabling contact between rRNA and an invariant ‘A’ nucleotide (Easton et al. 2009; Quade et al. 2015). All start codons were the canonical AUG. About 30% of the sequences included a well-documented additional subdomain IIId2 whose function is unclear (Li et al. 2024), but it may be more important in *Picornaviridae* IRESs than in *Flaviviridae* IRESs (Willcocks et al. 2017; Hashem et al. 2013b). Consistent with the literature, we found IIIabc to be variable (Asnani et al. 2015; Li et al. 2024), and in about 30% of our examples one or more of these subdomains was missing. Since many Type IV IRESs contain additional domains or lack one or more domains with unknown functional implications, using a single Type IV example to represent the entire class is likely misleading.

In contrast to the highly conserved core of domain III, domain II lacks a similar anchoring structure to help align variable regions including sequences with potentially unprecedented deletions and insertions, reducing the robustness of its alignment. The predicted structure of domain II is also quite variable, with 10% of the sequences having expansions compared to the HCV IRES. Potential implications of this variability are discussed in a later section.

In addition to the structural heterogeneity, there is a variety of host species for Type IV IRES containing viruses (**Fig. 1D**). Most hosts were mammals (mammalia) or birds (aves), with some from reptile (reptilia), amphibian (amphibia), and fish hosts (actinopteri). We found one example in a virus with a tick host (Amblyomma javanense), but probably the virus was in blood taken from an unknown host, rather than infecting the tick. Between vertebrates, ribosomal rRNA and protein as well as other translational machinery is highly conserved (**Fig. S1**), suggesting that these IRESs can function in diverse, and possibly multiple, host species (Brito Querido et al. 2024; Kyritsis et al. 2020; Zhao et al. 2013; Jubin et al. 2000).

### Validation and refinement of IRES alignment using chemical probing

Many putative Type IV IRESs have insertions or deletions in domains II and III. To confirm these predicted structures and better inform our alignment, we used selective 2′-hydroxyl acylation analyzed by primer extension and mutational profiling (SHAPE-MaP) (Siegfried et al. 2014; Smola et al. 2015) to biochemically validate IRES secondary structures of IRES RNA representatives from each cluster: Cluster 1 – hepatovirus C (HepV-C), Cluster 2 – limnipivirus C (LimV-C), Cluster 3 – megrivirus E (MeV-E).

SHAPE-MaP data support the IRES core alignment (IIIdef) for all three IRESs and for IIIabc for HepV-C and LimV-C (**Fig. 2A&B**). For MeV-E, we used our chemical probing data and the RNAprobing Webserver to predict the structure of IIIabc, revealing a significantly extended IIIa and IIIb, consistent with previous literature (**Fig. 2C**) (Hofacker 2003; Asnani et al. 2015). Twenty other IRESs contained a similar expansion. In domain II, the alignment predicted that HepV-C and LimV-C IRESs could have large insertions of flexible sequence. Using the SHAPE data with the RNAprobing Webserver to adjust the domain II secondary structure produced a model of the HepV-C IRES domain II secondary structure that includes an additional stem-loop element (**Fig. S2A**) (Hofacker 2003). Likewise, the LimV-C IRES domain II showed deviations from the alignment-derived consensus model, comprising two stem-loops (**Fig. S2B**). It is unclear if one or both together functionally replace domain II. In contrast with these two, the alignment’s predicted secondary structure of the MeV-E IRES domain II agrees well with the chemical probing and is similar to the HCV IRES (**Fig. S2C**).

**Figure 2.**
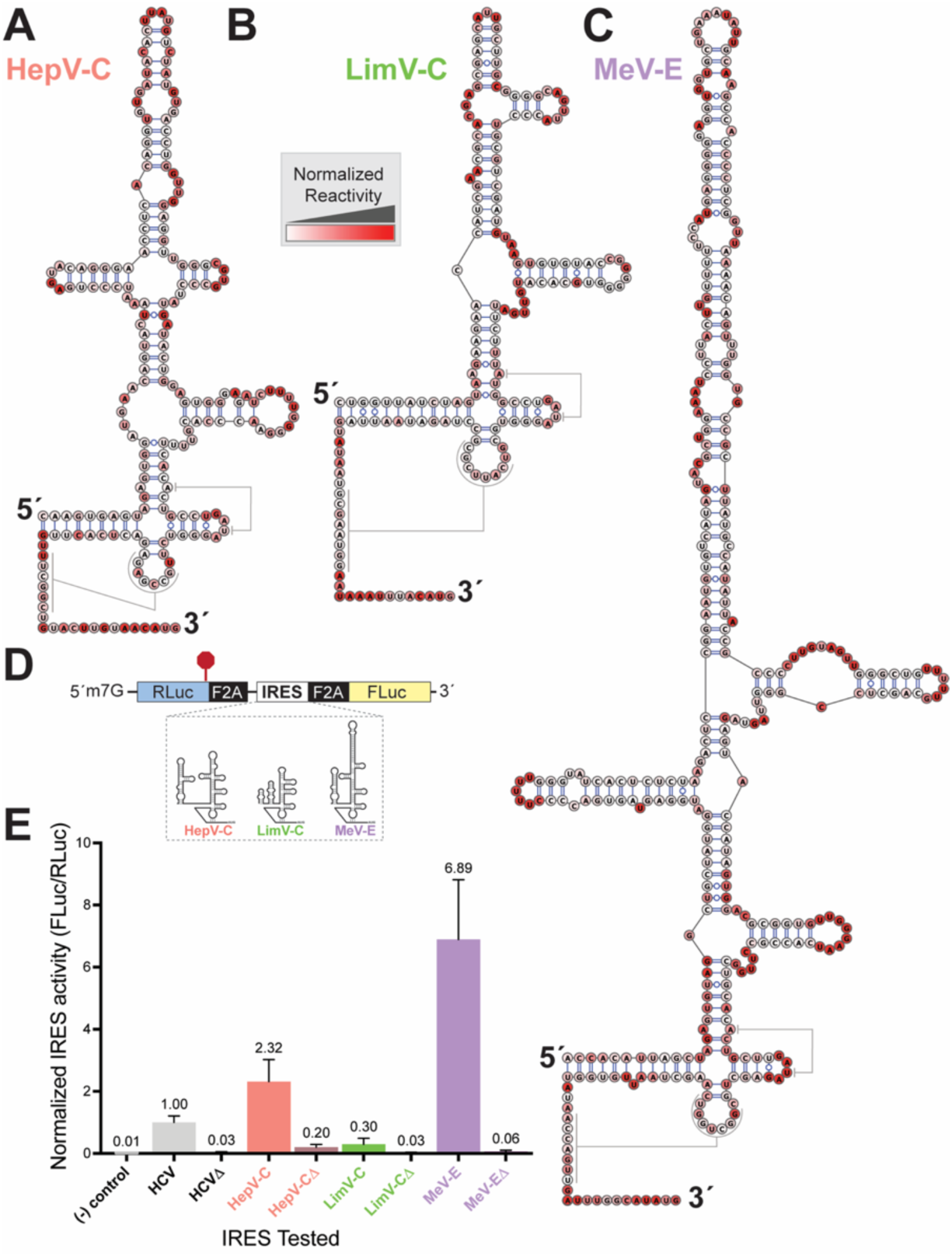
Validation of structure and functional tests. **(A-C)** Chemical probing results mapped onto domain III secondary structures. The color of each nucleotide indicates how reactive it was to the probing agent. **(A)** hepatovirus C (NC_038313; nt 324-530), **(B)** limnipivirus C (NC_039212; nt 327-504), and **(C)** megrivirus E (NC_039004; nt 376-720). Hepatovirus C and limnipivirus C results are mapped on to alignment-predicted secondary structures. Megrivirus E results are mapped on to a secondary structure where IIIabc (408-616) was predicted using the RNAprobing webserver with SHAPE results. Results show domain III up to the start codon. **(E)** In vitro translation assays of IRES activity using dual luciferase constructs. RLuc: Renilla luciferase (RLuc), FLuc: Firefly luciferase, F2A: peptide bond-skipping sequences, red octagon: stop codon. Cartoon secondary structures of the three IRESs tested are shown. **(D)** Graph contains the results of translation assays of both wild-type and mutant (Δ) IRES sequences, normalized to wild type HCV IRES.

Overall, these results suggest that our alignment accurately predicts domain III structure but is limited by variation in both shorter and longer domain II sequences. Using experimental data, we were able to improve the alignment in areas with uncertain structure. These biochemical approaches also confirm previously predicted – but untested – IRES structures (Asnani et al. 2015).

### Clustering Type IV IRESs by secondary structural patterns

The large size and vast structural heterogeneity among Type IV IRES sequences demanded a more rigorous approach than relying on “by eye” comparison. We therefore employed a clustering algorithm to group IRESs based on structural features, focusing on domain III.

For each putative Type IV IRES we quantified (1) the number of base pairs in each domain III subdomain stem, (2) the number of nucleotides in their apical loops, (3) the number of base pairs in the IIIf pseudoknot, (4) the number of nucleotides from the end of the pseudoknot to the start codon, and (5) the total length of domain III. We then applied the CLARA (Clustering Large Applications) algorithm, a partitioning method optimized for efficiently handling large datasets. CLARA clustering was applied directly on the full feature set without prior dimensionality reduction yielded 3 clusters (**Fig. S3; Fig. S4**), defined largely by the lengths of domain III and IIIb (**Fig. 3**).

**Figure 3.**
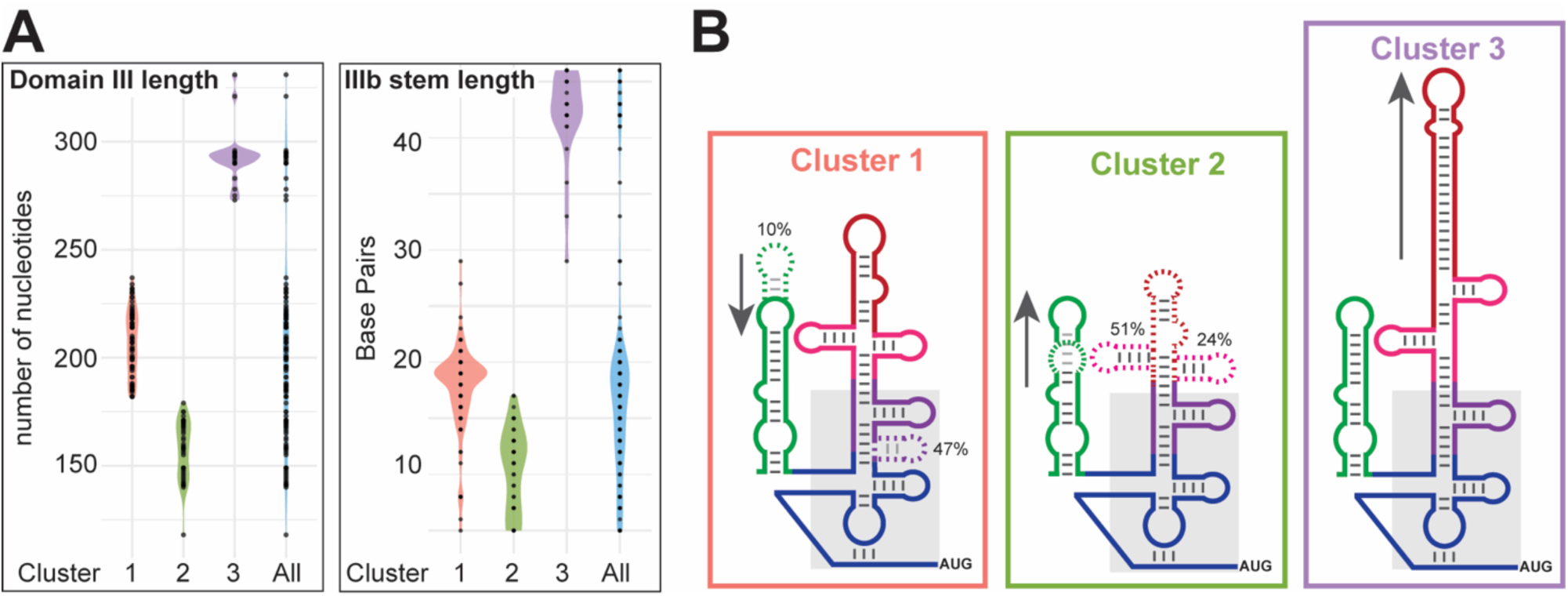
Cluster analysis of Type IV IRESs. **(A)** Violin plots of the distribution of cluster-defining structural features for each cluster (1-3) and all IRESs together (All). Total domain III length as measured in nucleotides (left) and subdomain IIIb stem length as measured in base pairs (right) are included. **(B)** Cartoon schematics of IRES clusters. “Cluster 1” IRESs maintain all the same subdomains as the HCV IRES. They can also include an additional domain, most often IIId2. “Cluster 2” IRESs have truncated subdomains IIIabc, if not lacking them entirely. “Cluster 3” IRESs have elongated IIIa and IIIb. Arrows and dashed regions indicate areas of distinct variability.

“Cluster 1” includes the prototype HCV IRES (**Fig. 3**). These IRESs have an average domain III length of 207 nucleotides with a range of 182 - 237, and ∼50% contain domain IIId2. The average size of IIIb is 12 base pairs with a range of 0 - 24, and IIIa and IIIc loop sequences are conserved (**Fig. S5A**). This cluster includes Type IV IRES examples in the viral families *Flaviviridae*, *Picornaviridae*, *Caliciviradae* and *Picobirnaviridae* (**Table S2**). Hosts species are limited to mammals and birds with two examples with unknown hosts. Additionally, there is one example found in an insect, but it is unclear if the insect is the true host of this virus (**Table S2**).

“Cluster 2” are smaller and lack part or all IIIabc (**Fig. 3**). The average domain III length is 157 nucleotides with a range of 118 - 179, and these IRESs are on average ∼50 nucleotides shorter than those in Cluster 1. When present, IIIb is much smaller than those in Cluster 1, with an average stem length of 5 base pairs with a range of 0 - 12. Cluster 2 largely lacks sequence conservation but maintains the predicted tertiary interactions involving IIIf and IIIe (**Fig. S5B**). Most of the IRESs in this cluster are found in *Picornaviridae* and unclassified viruses (**Table S2**). Cluster 2 has the most diverse array of host species, including fish, amphibians, birds, reptiles, and mammals.

“Cluster 3” IRESs are the largest by IRES size, with a significantly expanded IIIb (**Fig. 3**). The average domain III length is 294 nucleotides, with a range of 273 - 331. Generally, IRESs in Cluster 3 are about 50 nucleotides longer than those in Cluster 1, with an average IIIb length of 37 base pairs ranging from 24 - 41. This size of IIIb is about 25 base pairs longer than those in Cluster 1. Interestingly, the apical loop of IIIb is highly conserved within this cluster (**Fig. S5C**). This ‘figure-8’ motif has been described previously, but it’s not clear if or how it impacts IRES function (Asnani et al. 2015; Li et al. 2024). IIIa is also longer, with an average of 10 base pairs compared to the Cluster 1 average of 6 base pairs (**Fig. S5C**). Of the 22 IRESs in this group, all are in *Picornaviridae*, with most in the megrivirus genus, one in each anativirus and tropivirus, and several being unclassified. All the viruses with Cluster 3 IRESs have birds as the identified host species expect for one, which has gecko as the host species (**Table S2**). Additionally, we noticed that there was an IRES example in Cluster 1 (KT880670.1) with IIIb sequence conservation and structure generally consistent with Cluster 3 IRESs. The IRES lacks IIIa and has a shorter IIIb than most examples in Cluster 3, which likely placed it in Cluster 1 as our clustering analysis was sequence agnostic.

### Functional variation among Type IV IRESs

To correlate variation in Type IV IRES clusters with function, we performed *in vitro* translation assays with a dual-luciferase reporters containing an upstream, cap-driven *Renilla* luciferase (Rluc) as an internal control and a downstream, IRES-driven Firefly luciferase (Fluc) (**Fig. 2D**). The downstream Fluc sequence is out of frame with the Rluc to prevent ribosome readthrough. Peptide bond-skipping sequences (F2A) prevent artifacts due to inserted sequences that are initially translated but then liberated from the mature luciferase proteins. A reporter RNA lacking an IRES served as negative control while a reporter RNA where the HCV IRES drives Fluc expression is the positive control. IRES activity was normalized to the positive control.

We again tested the same representative IRES from each of the three clusters. In addition to wild-type IRES sequences, we also tested IRES RNAs carrying a mutation in the loop of IIId: GGG → CCC, denoted as Δ. This mutation decreases HCV IRES activity by disrupting a key interaction with the 40S subunit (Kieft et al. 2001). IRESs with this mutation were included to test the prediction that all use this interaction as a critical part of their mechanism.

For our assay system, we used rabbit reticulocyte lysate. The HepV-C IRES initiated translation at a level very similar to the HCV IRES, consistent with a similar secondary structure in domain III (**Fig. 2E**). In contrast, the LimV-C IRES had lower translation initiation efficiency, ∼30% of that shown by the HCV IRES. In fact, we were surprised that the LimV-C demonstrated any IRES activity at all, as it lacks IIIa and has a truncated IIIb. Finally, MeV-E IRES had ∼7 fold higher activity than the HCV IRES. All mutant IRES had dramatically reduced activity relative to their wild type counterpart, thus all likely interact with the ribosome using the conserved core domain.

### Potential mechanistic implications of structural variation

We used an existing structure of the HCV IRES bound to the ribosome (Quade et al. 2015), and the known roles of different domains, to examine how structural variation in Type IV IRESs might alter these key interactions and thus IRES mechanism. First, in HCV domain II is flexible, adopting multiple stable configurations with additionally freedom conferred by a single-stranded RNA linker proceeding domain III (Boerneke et al. 2014; Lukavsky et al. 2003). Domain II contacts the 40S subunit’s E site to help position the coding RNA in the decoding groove and facilitate later stages of 80S assembly and transition to elongation (Honda et al. 1999; Locker et al. 2007; Quade et al. 2015; Filbin and Kieft 2011; Filbin et al. 2013). An expanded domain II could make the same contacts but perhaps with more intervening structure. A larger bifurcated domain II could make additional contacts with the ribosome or perhaps interact with another factor (**Fig. 4**). For a shorter domain II, it is not clear how it could reach into the E site, as in the HCV and CSFV IRES the domain must be long enough to “bend” into position (Locker et al. 2007; Honda et al. 1999). Such IRESs, and those with even more highly divergent domain II regions, may use alternate contacts or mechanisms; this remains to be explored.

**Figure 4.**
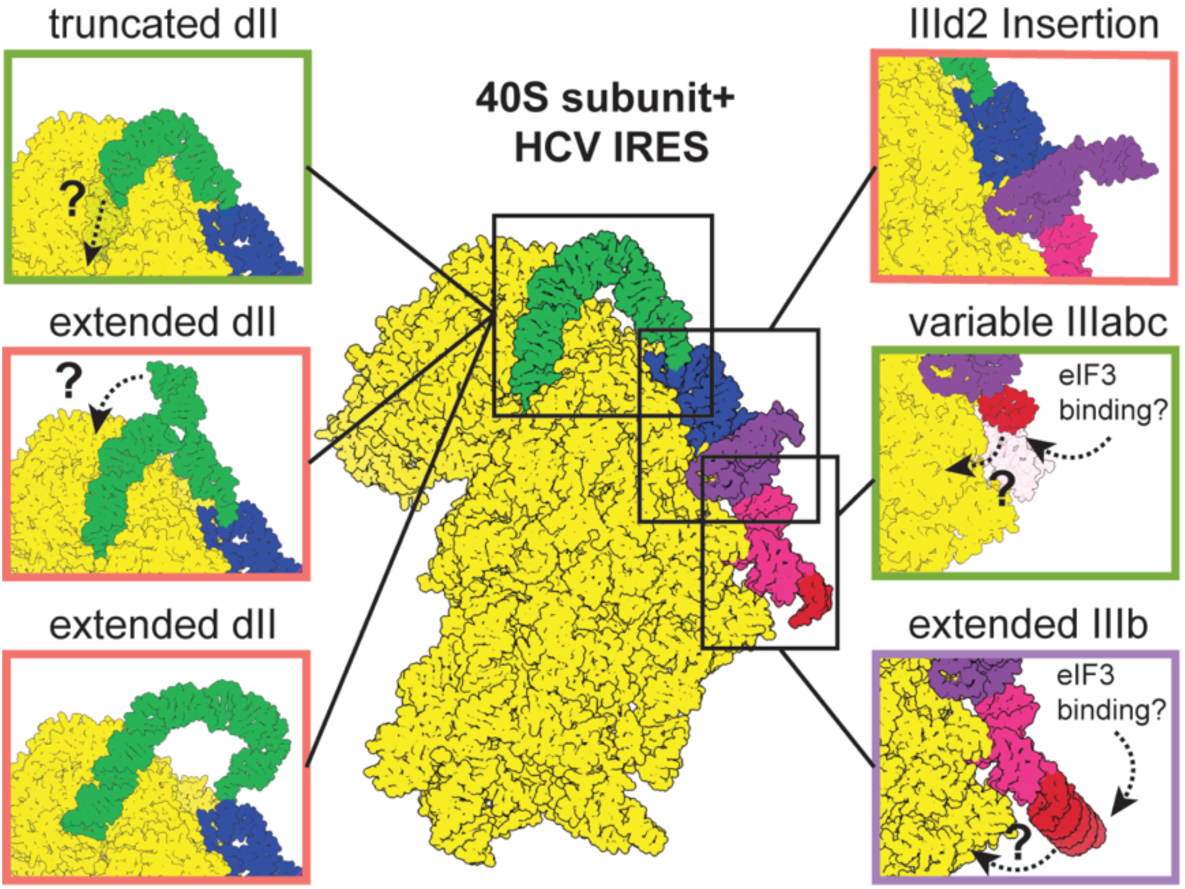
Modeling implications of Type IV IRES structural diversity. Cryo-EM structure of the HCV IRES (colored by domain as in Fig. 1A) bound to with mammalian 40S subunit in yellow (PDB: 5A2Q). Regions of variability within the Type IV IRESs are boxed, and individual panels show possible implications of this diversity.

Second, in the HCV IRES domain IIIabc binds initiation factor eIF3 and the 40S subunit (Pestova et al. 1998; Sizova et al. 1998; Kieft et al. 2001; Sun et al. 2013), thus for IRESs with an HCV-like domain IIIabc (i.e. Cluster 1), these interactions are likely maintained. However, because the nucleotide- and residue-resolution details the interaction are unknown, it is challenging to predict how IRES variability affects eIF3 binding (Sun et al. 2013; Hashem et al. 2013b). For those lacking much of IIIabc (i.e. Cluster 2) (**Fig. S5B**), critical interactions are likely lost (Hashem et al. 2013b; Sun et al. 2013) (**Fig. 5**), consistent with the effect of IIIabc HCV mutations on the HCV IRES (Kieft et al. 2001). IRESs with extended IIIabc (i.e. Cluster 3) (**Fig. S4C**) might bind eIF3 using more of its subunits, could recruit additional unknown proteins, or make additional contacts with the ribosome (**Fig. 4**). While there is much sequence and structural variability in IIIabc, the proteins theorized to contact these domains are conserved across vertebrates (**Fig. S1B-D**).

Third, an additional source of variability is the presence or absence of IIId2 (**Fig. 5; Fig. S5A**). IRESs that contain this domain are mostly in Flaviviridae, where it is functionally important (Willcocks et al. 2017). However, a few picornaviruses (Senecavirus, Parechovirus, and Tropivirus) include this domain. Within Senecaviruses, IIId2 is not required for IRES activity, but deletion of this domain does affect viral growth and survival (Willcocks et al. 2017).

Finally six IRESs, from the Sapelovirus and Ludopivirus genera (NC_004451, AY064715, JX627574, JX627573, AY064720, NC_040684), also contain a small stem-loop between IIIa and IIIb, referred to as IIIa2 (Oberste et al. 2003; Boros et al. 2018). Other studies have identified several Kobuviruses with an additional domain between IIIa and IIIb while lacking IIIc (Reuter et al. 2009; Asnani et al. 2015). Our alignment assigned these three stems sequentially, as IIIabc. The previous study may have annotated IIIa2 to avoid an unusually long IIIc, which would have seemed inconsistent with existing knowledge at the time. However, based on our alignment and biochemical data, an elongated IIIc appears plausible. Notably, the role of these additional domains in IRES function and viral survival has not been studied.

### Summary

We have cataloged the structural diversity among Type IV IRESs and identified a new example in viral clades not formerly known to have them. These IRESs share a core functional domain but show diversity in functionally important peripheral domains, which confers variability in function and perhaps in mechanism. This variability, combined with potentially broad species tropism and reduced reliance on initiation factors, makes them useful tools both for molecular biology and medical science. Future work to fully understand and characterize these diverse IRESs will be critical for realizing their potential.

## METHODS

### Alignment and homology search

A preliminary Type IV IRES alignment was created using six unique IRESs with sequences from the National Center for Biotechnology Information (NCBI) sequence database (accession numbers: NC_004102, NC_002657, MZ334483, NC_001655, NC_004451, HQ020378). Existing alignments of Type IV IRESs (Rfam IDs: 00061, 00209) were obtained from the Rfam database, and their secondary structure models were combined (Ontiveros-Palacios et al. 2025). The sequences were manually aligned in Stockholm format based on primary features (IIId apical ‘GGG’ motif, start codon), secondary structural features (domain II, subdomains IIIa-f), and tertiary structural features (IIIf pseudoknot and IIIe long-range base pair). This seed alignment was used to query a RefSeq viral database (01/22/2019), using Infernal v1.1 (Nawrocki and Eddy 2013). A total of 176 unique sequences were returned from the search. Thirteen sequences were dropped from the alignment due to degenerate sequence or undeterminable secondary structure features (**Table 1**). The alignment of the remining 163 sequences was manually adjusted based on chemical probing data and RNAVienna predictions (Hofacker 2003). The resulting alignment was used to generate a consensus model calculated using R-Scape, graphically depicted in R2R, and adjusted in Adobe Illustrator (**Fig. 1C**; Rivas 2020; Weinberg and Breaker 2011).

### Dataset generation and clustering analysis

The secondary structure characteristics of each IRES were analyzed using a custom Python (v3.9.6) script. The Stockholm file format and IRES feature coordinates were maintained during analysis using the Biopython (v1.81) AlignIO package. The script identifies and quantifies stem-loop structures (II, IIIa–f), the pseudoknot (PK) based on consensus secondary structure annotations. It calculates stem and loop lengths, 5′ UTR length, domain III size, distance from the PK to the start codon, and sequence leading up to domain II. The script also accounts for sequence degeneracy. All measurements were reported in a CSV file for downstream analysis. Custom python scrips are available through a Github public repository (https://github.com/segarka/TypeIV_IRESs).

Subsequent clustering analysis was performed in R studio (v2023.09.1+494). Clustering was performed using the cluster package (v2.1.8.1; Maechler et al. 2025). Analysis was applied to the full feature set without initial dimensionality reduction. The optimal number of clusters (k = 3) was selected by eye using the elbow method, which identifies a point of diminishing returns in within-cluster sum of squares (factoextra v1.0.7; Kassambara and Mundt 2020). We initially tested two clustering algorithms: Partitioning Around Medoids (PAM) and Clustering Large Applications (CLARA). Both are classical and valid clustering methods (Kaufman and Rousseeuw 1990). However, the CLARA algorithm produced clusters that were more biologically interpretable and thus used for sub-setting the larger alignment. Violin plots were generated using ggplot2 (v3.5.2; Wickham et al. 2019).

### Taxonomy and phylogeny

Viral and host taxonomy were mined from the NCBI database using taxonomizr (v0.11.1). For viral phylogenic tree construction, representatives of each viral class as well as all unassigned viruses were included. The entire genome of the viruses were aligned with ClustalW using the msa package (v1.34.0; Bodenhofer et al. 2015). The phylogenic tree was constructed and visualized using phylotools (v0.2.2) and edited in Adobe Illustrator.

### RNA Preparation for translation assays

The pSGDlucV3.0 plasmid (vector) and gBlocks (inserts) were prepared by digesting with BglII in NEBuffer r3.1 (NEB) at 37°C for 2 hours. PspXI and rCutsmart™ Buffer (NEB) were added to double the volume of the initial reaction and incubated at 37°C for 2 hours. The vector was purified via agarose gel purification with the Wizard SV Gel and PCR Clean-up System (Promega) and eluted in water. Inserts were the purified using the same kit, eluted in water. Both vector and inserts were quantified using Nanodrop Spectrofluorometer (Thermo Scientific).

Inserts were incorporated into the vector via ligation. The ligation reaction consisted of inserts and vector in a 3:1 ratio, Quick ligase (NEB), 2X Quick ligase buffer (NEB). The reaction was incubated at room temperature for 10 minutes and half the reaction was transformed into NEB stable cells and plated on +Amp LB plates. Colonies were prepped and plasmids were sent for full plasmid sequencing before use. IRES Dual-Luciferase constructs (DLuc) were amplified via PCR and cleaned up using the Nucleospin PCR clean-up kit (Macherey-Nagel). PCR products were size verified on a 1% agarose gel. PCR produced were used as templates for the T7 HiScribe Transcription kit (NEB) and co-transcriptionally capped by including ARCA (NEB) in the reaction according to the kit instructions. RNA was purified using the Monarch RNA Cleanup kit (NEB) and eluted in water.

#### RNA preparation for chemical probing

IRES sequences were amplified via PCR from plasmids. Forward and reverse primers were specific to each IRES construct and encapsulated the entirety of the IRES coordinates and sequence for 15 N-terminal codons, including the initiation methionine codon. PCR products were size verified on a 1% agarose gel. PCR products were used as templates for large-scale transcription reactions which included the following components at noted final concentration: ATP (6mM), GTP (6mM), UTP (6mM), CTP (6mM), MgCl2 (42 mM), 10X transcription buffer (1X; 30 mM Tris-HCl (pH 8.0), 10 mM DTT, 0.1% spermidine, 0.1% Triton X-100), T7 RNA polymerase, and RNase Inhibitor. Transcription reactions were incubated overnight at 37°C. RNA was purified via ethanol precipitation followed by gel purification on a denaturing 6% acrylamide gel. RNA was eluted from acrylamide by overnight incubation at 4°C in 200 mM NaOAc (pH 5.2). RNA was then concentrated via ethanol precipitation and resuspended in water.

All RNA was quantified using Nanodrop Spectrofluorometer (Thermo Scientific). Quality was assessed using 260/280 and 260/230 ratios as well as visually on 10% acrylamide gels via ethidium bromide staining.

### SHAPE-MaP

The SHAPE-MaP protocol was followed as outlined in this manuscript (Smola et al. 2015). Briefly, 1 µg of IRES RNA in 12 µL of DEPC treated water was heat re-folded by incubation at 95°C for 1 minute and cooled to room temperature for 5 minutes. After cooling, 0.5 µg of RNA was labeled with 10 mM 1-methyl-7-nitroisatoic anhydride (Millipore Sigma) or the control, DMSO, for 75 seconds at 37°C. Modified and control samples were purified and concentrated using the RNeasy Micro Kit (Qiagen), eluting in 12 µL of DEPC water.

Following modification and purification, modified/control RNA was incubated with 200 ng/µL Random Primer 9 mix (NEB) and 1 µL of custom reverse primer specific to the IRES at 65°C for 5 minutes and then cooled on ice. To each reaction, 8 µL of MaP buffer (125mM Tris pH 8.0, 187.5mM KCl, 15 mM MnCl_2_, 25 mM DTT, 1.23 mM equimolar dNTP mix, and 200U Superscript II) was added to the reaction and incubated at 25°C for 2 minutes, 25 °C for 10 minutes, 42°C for 3 hours, 70°C for 15 minutes, and cooled to 4°C. The RNA/cDNA hybrids were then purified on the G-25 Microspin columns (Cytiva) spin column following the manufacturers protocol after slight modification. Before purification the columns were washed 3x with 500ul of DI water to prevent carryover DNA contamination from the storage buffer. Second strand synthesis of the cDNA was carried out using the NEBNext Ultra II Non-Directional RNA Second Strand Synthesis Module (NEB) following the manufacturer’s instructions. The resultant cDNA was purified using the DNA Clean & Concentrator-5 kit (Zymo) eluting in 10 ul of DEPC-treated H_2_O. The resultant dsDNA pool was quantified fluorometrically with a Qubit 3 Fluorometer (Thermo) using the Qubit dsDNA HS Assay Kit (Thermo). Next generation sequencing library prep was performed with the Nextera XT DNA library preparation kit (Illumina) using the Nextera XT Index Kit v2 Set B (Illumina). Samples were normalized and pooled together using a combination of the Qubit and the 4200 TapeStation System (Agilent Technologies) with the High Sensitivity D5000 Screen Tape (Agilent Technologies). Samples were sent to Novogene for sequencing on the NovaSeq6000 asking for 10million paired end reads for each sample.

For SHAPE-Map analysis, raw sequencing reads were first visualized with FastQC (v0.11.9) to assess the quality of the data. The raw reads were then trimmed using Trimmomatic (v040), to remove any residual Illumina adaptors. The processed reads were then analyzed with Shapemapper2 with the default parameters and a minimum read depth of 5000 reads. The resultant data was analyzed using a custom in house pipeline of Python (v3.9.10) and R (v4.2.2) scripts. Average reactivities of three biological replicates per construct were then plotted onto the proposed secondary structure using Varna (v3.93) (Darty et al. 2009) and manually edited in Adobe Illustrator.

### In vitro translation assays

RNA was prepared in 4.5 µL volume with 1 µg of RNA and 33mM HEPES (pH 7.5). RNA was heat re-folded by incubating at 95°C for 2 min and cooling back to room temperature for 5 minutes. MgCl_2_ was then added to a concentration of 10 mM to yield a final reaction volume of 5 µL. The final translation reaction mixture included the RNA mixture described above and components from the Rabbit Reticulocyte Lysate system (Promega): rabbit reticulocyte lysate (35 µL), amino acid mixture minus cysteine (final concentration 16.7 μM), amino acid mixture minus methionine (final concentration 16.7 μM), amino acid mixture minus leucine (final concentration 16.7 μM), KOAc (to final concentration of 150 mM), and water to a final reaction volume of 50 µL. Reactions were incubated at 30°C for 3 hours.

After incubation, reactions volume were adjusted to 250 µL using 1X Passive Lysis Buffer (Promega). The reaction was split into 2 wells of 100 µL each. Luciferase activity was measured using Dual-Luciferase Reporter Assay System (Promega) with the GlowMax-Multi+ Detection System (Promega) with the pre-programed Dual-injector, Dual Luciferase protocol. Data were analyzed in Excel, plotted using GraphPad prism (v10.3.0), and labeled using Adobe Illustrator.

## ACKNOWLEDGEMENTS

Thank you to current and former Kieft Lab members for thoughtful discussion and technical assistance. pSGDlucV3.0 was a gift from John Atkins (Addgene plasmid 119760). This work was supported by NIH grants 1F31AI186389-01 (K.E.S), T32GM136444 (K.E.S), R35GM118070 (to J.S.K), and a McKnight Foundation Award (to J.S.K). M.E.S was supported by a Jane Coffin Childs postdoctoral fellowship.

