## Supplemental Figures and Tables for "Distribution and structural diversity of Type IV internal ribosome entry sites"

**<u>Supplemental information contents:</u>**

**Supplemental Figure S1:** Conservation of key rRNA and translation initiation factor sequences in vertebrates.

**Supplemental Figure S2:** Chemical probing of domain II.

**Supplemental Figure S3.** Sub-grouping of structurally similar type IV IRESs using cluster analysis.

**Supplemental Figure S4.** Additional data on sub-groups secondary structural features.

**Supplemental Figure S5.** Cluster consensus/covariation models.

**Supplemental Figure S6.** Additional functional data on putative IRESs in rabbit reticulocyte lysate.

**Supplemental Figure S7.** Functional data on putative IRESs in a human *in vitro* translation system.

**Supplemental Table S1:** Sequences discarded from the covariation and clustering analysis.

**Supplemental Table S2:** Taxonomic information on viral family and host class between Clusters.

**Supplemental Table S3:** DNA oligonucleotide sequences used in this study.

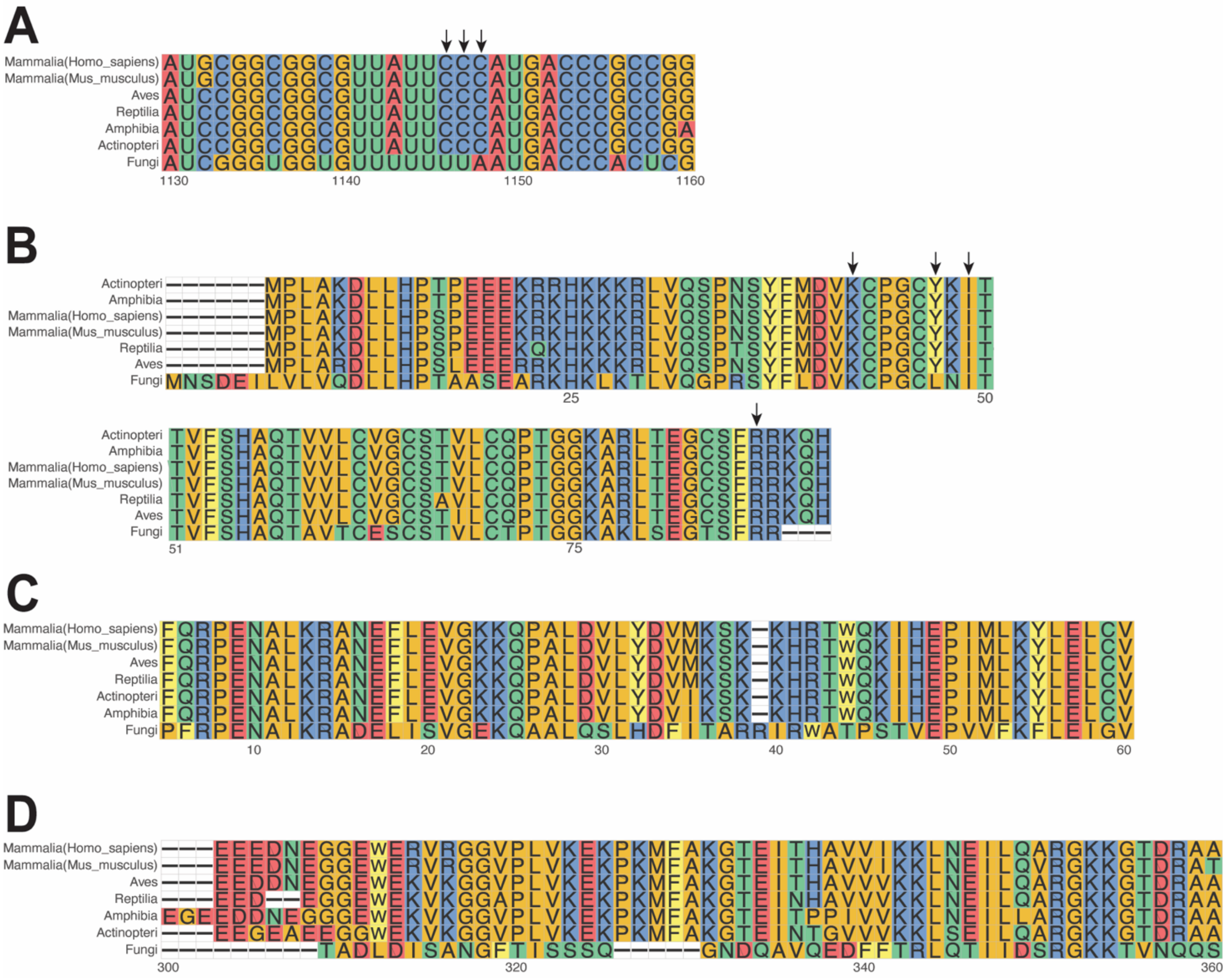


**Figure S1. Conservation of key rRNA and translation initiation factor sequences in vertebrates.** (*A*) Alignment of 18S rRNA sequences focused on expansion segment 7 across representative vertebrate examples. Arrows show conserved ‘CCC’ motif. Nucleotides are colored by identity. (*B*) Alignment of ribosomal protein eS27 across representative vertebrate examples. Arrows show conserved residues that were implicated in HCV IRES interaction (*Quade. et al. 2015*). (*C*) Alignment of eIF3A N-terminal region, containing helix-loop-helix motif that is suggested to directly bind the HCV IRES (Sun et al. 2013). (*D*) Alignment of eIF3C helix-loop-helix motif that is suggested to directly bind the HCV IRES (Sun et al. 2013).

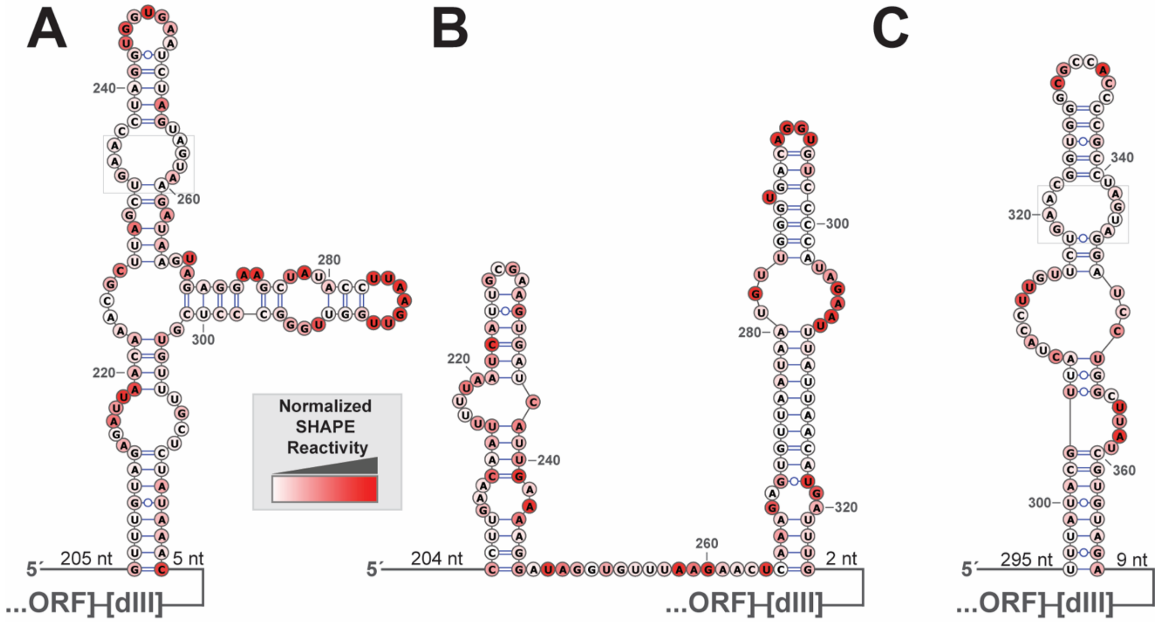


**Figure S2.** Chemical probing of domain II. Chemical probing results mapped onto secondary structure of domain II from different IRESs. The color of each nucleotide indicates how reactive it was to the probing agent (inset).**(A)** hepatovirus C (NC_038313:206-318), **(B)** limnipivirus C (NC_039212:205-324), and **(C)** megrivirus E (NC_039004:295-366). Domain II secondary structure for hepatovirus C and limnipivirus C was predicted with chemical probing data and sequence using the RNAprobing Webserver. Domain II secondary structure for megrivirus E was pulled directly from the alignment. The number of nucleotides upstream of domain II and between domain II and domain III are indicated. Loop E motifs are boxed in gray.

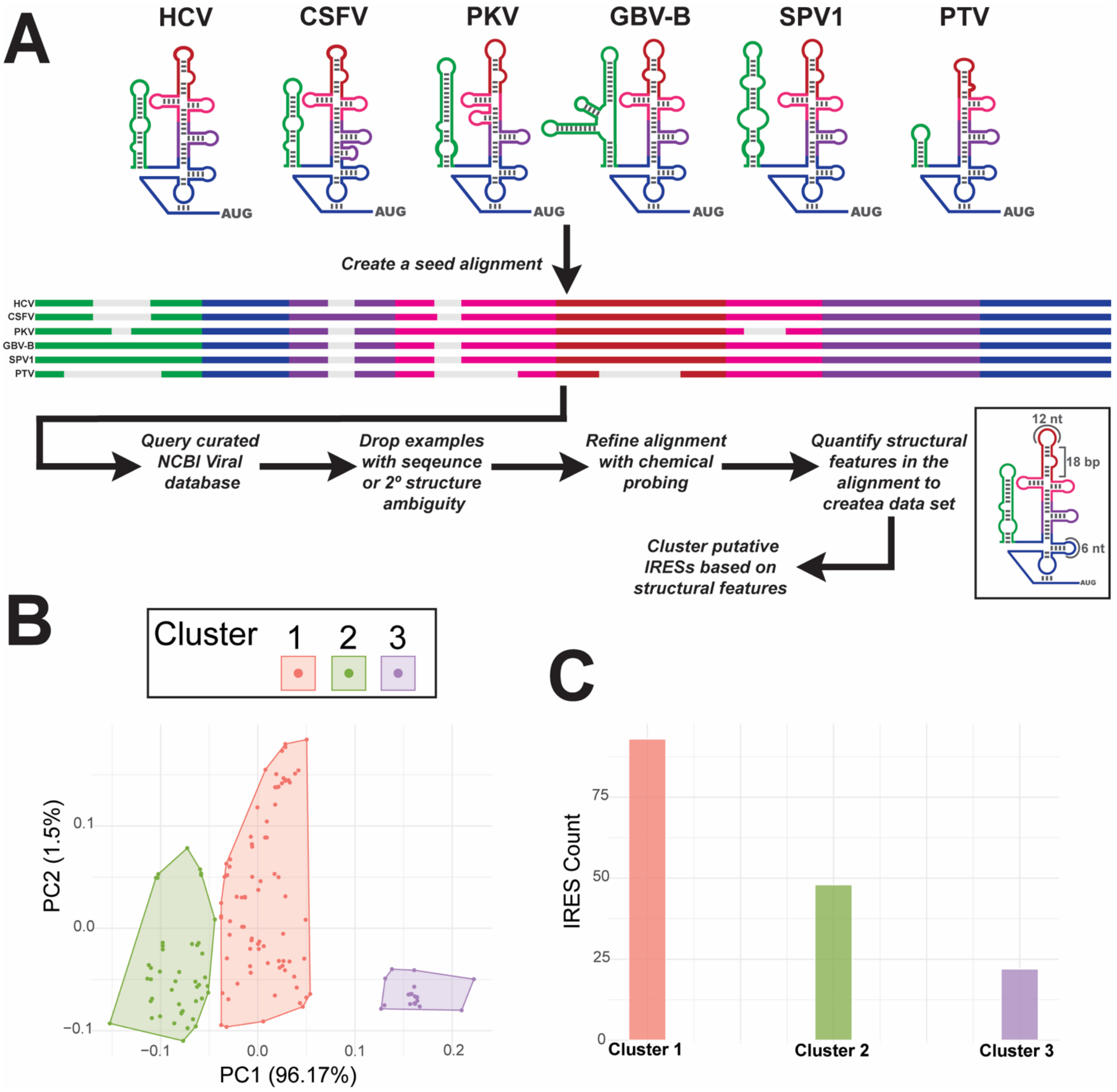


**Figure S3.** Sub-grouping of structurally similar type IV IRESs using cluster analysis. (*A*) Workflow showing how the data set for analysis was generated. Six type IV IRESs were aligned based on secondary structure and important sequence features to create a seed alignment. The seed was used to search a curated NCBI database, limited to roughly one example per viral species. Of the 176 hits, 13 were dropped due to structure or sequence ambiguity. Chemical probing data on a diverse selection of putative IRESs was used to further refine the alignment. Subdomain features (i.e. stems and loops) were quantified using custom python scripts, which counted the number of base pairs or nucleotides present in a feature for each IRES. This dataset detailing structural features for each IRES was then used for clustering analysis. (*B*) Principal component analysis (PCA) plot of clustering results. Data was clustered into 3 groups using the CLARA algorithm (detailed in Methods). The first two principal components (PC1 and PC2), computed via PCA, are shown on the x- and y-axes. Each point represents an individual data observation, and colors indicate the assigned cluster membership. Convex hulls enclose points belonging to the same cluster for visualization purposes. (*C*) Graphical representation of the number of IRESs are in each subgroup.

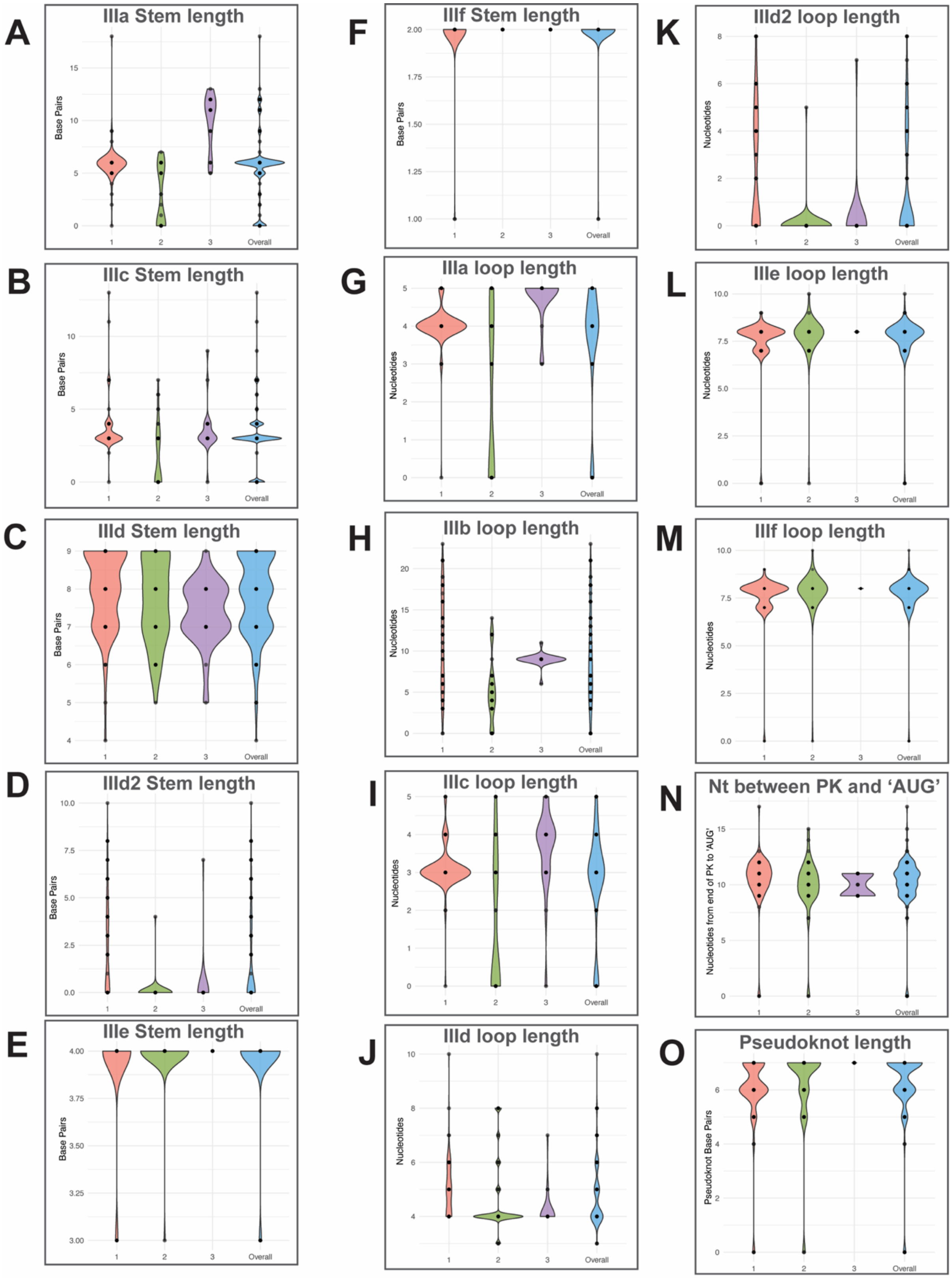


**Figure S4. Additional data on sub-groups secondary structural features.** Violin plots of the distribution of sub-domain stem lengths as measured in base pairs: (*A*) IIIa, (*B*) IIIc, (*C*) IIId, (*D*) IIId2, (*E*) IIIe, (*F*) IIIf. Violin plots of distribution of sub-domain loop lengths as measured in nucleotides: (*G*) IIIa, (*H*) IIIb, (*I*) IIIc, (*J*) IIId, (*K*) IIId2, (*L*) IIIe, (*M*) IIIf. (*N*) Violin plot of distribution of nucleotides between the end of the pseudoknot and the start codon (AUG). (*O*) Violin plot of distribution of pseudoknot length as measured in base pairs.

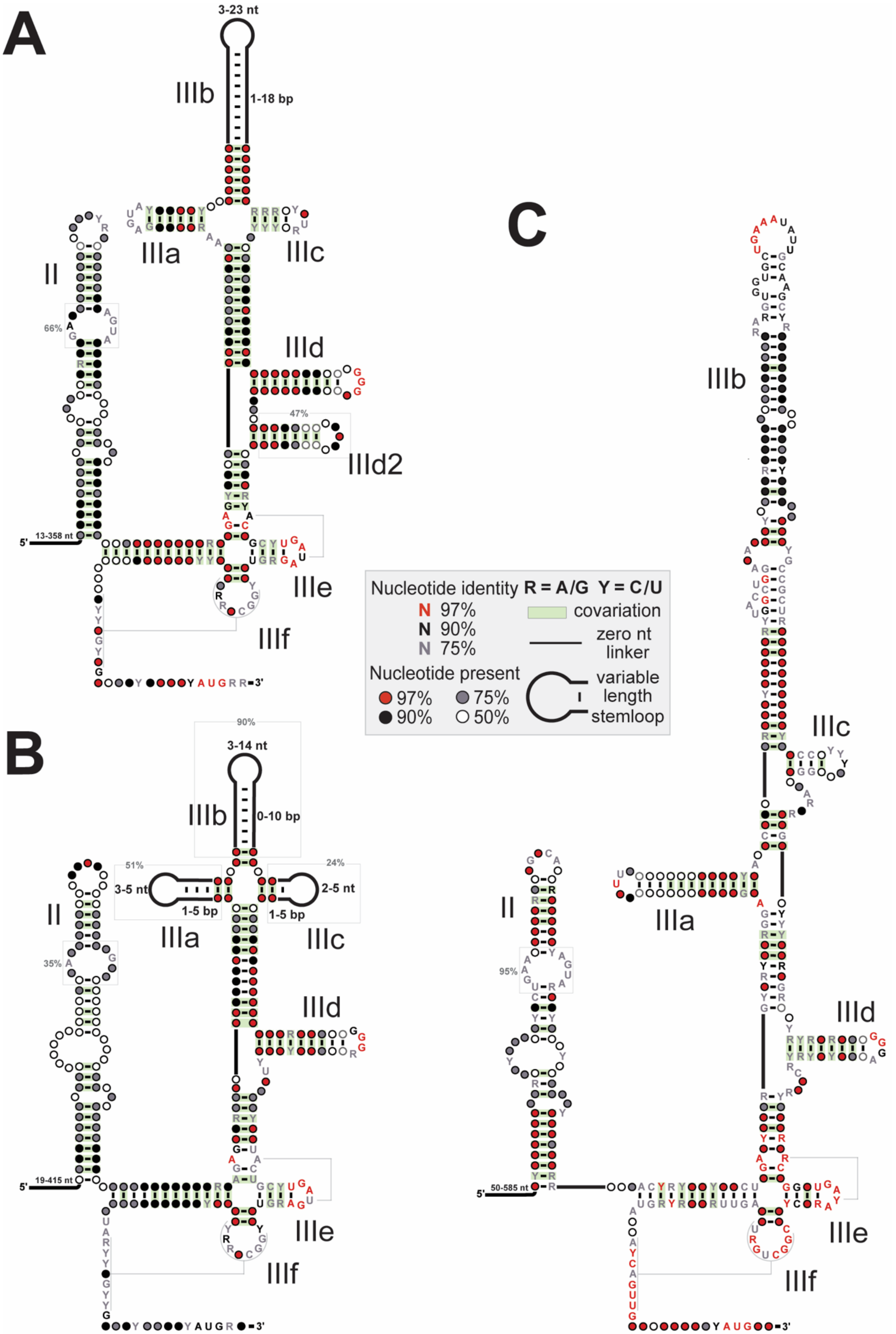


**Figure S5. Sequence and secondary structure consensus models for**“Cluster 1” **(A)**, “Cluster 2” IRESs **(B)** and “Cluster 3” IRESs **(C)**, detailing conserved structure, as determined by statistically significant covariation, and sequence features. Models generated using R-scape

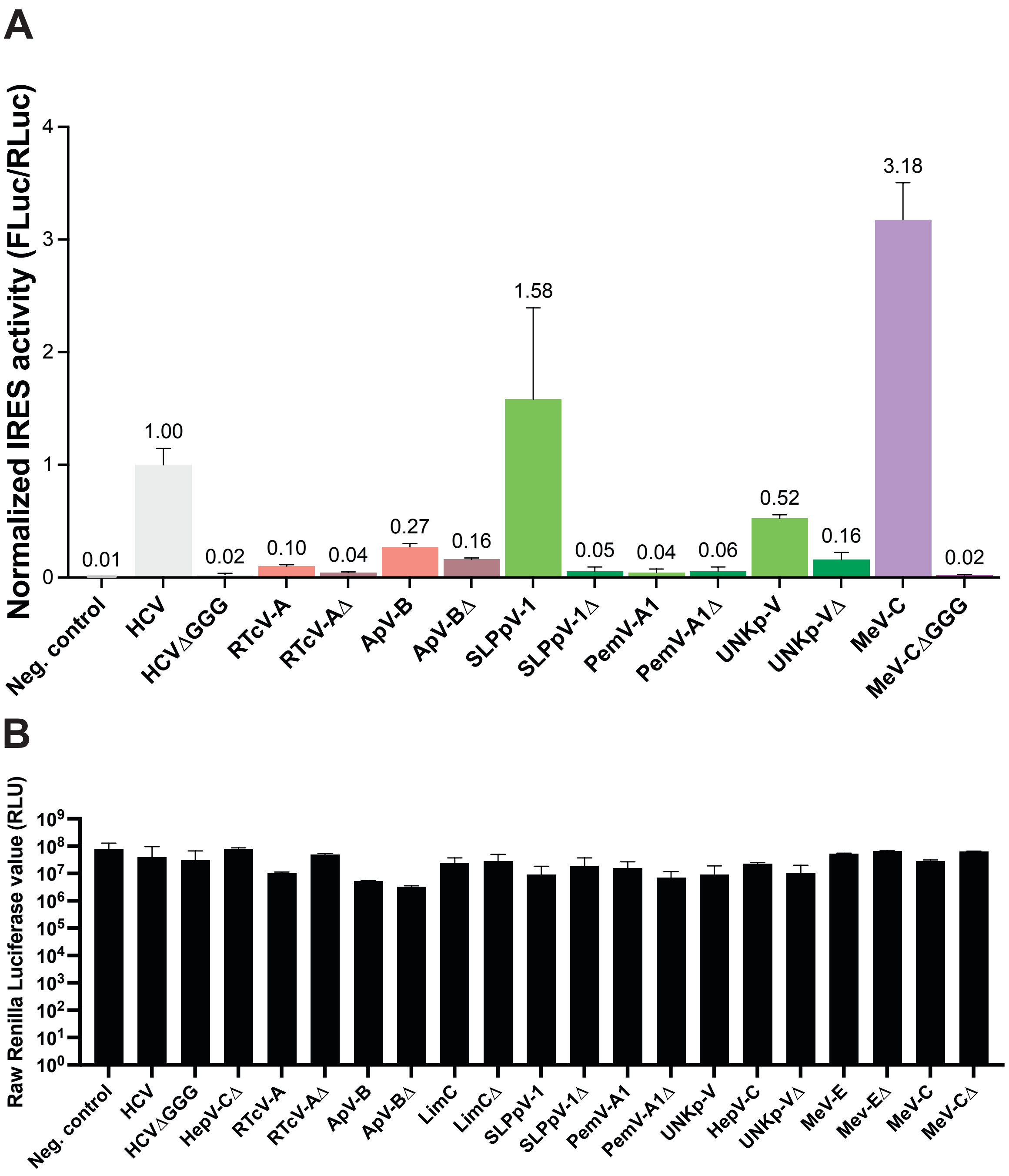


**Supplemental Figure S6.** Additional functional data on putative IRESs in rabbit reticulocyte lysate. In vitro translation assays of IRES activity using dual luciferase constructs in rabbit reticulocyte lysate. Dual luciferase constructs are as described in Figure 2. **(A)** The graph contains the results of translation assays of both wild-type and mutant (Δ) IRES sequences, normalized to wild type HCV IRES. The IRESs shown are as follows: Ruddy turnstone calicivirus A (RTcV-A, MH453861:200-581), avocet picornavirus B MW14 (ApV-B, MH453809:95-458), sapelo-like bat picornavirus 1 (SLPpV-1, HQ595341:16-350), Pemapivirus A1 (PemV-A1, MG600106:394-713), picornaviridae sp. isolate wftcra74pic2 (UNKp-V, MT138373:249-558), megrivirus C (MeV-C, HM751199:50-506). The color of the bars indicates the Cluster each IRES belongs to; pink for Cluster 1, green for cluster 2, purple for Cluster 3. **(B)** Raw *Renilla luciferase* values for each construct tested in the rabbit reticulocyte system. Virus names and genomic information of IRESs are as described above and in Figure 2.

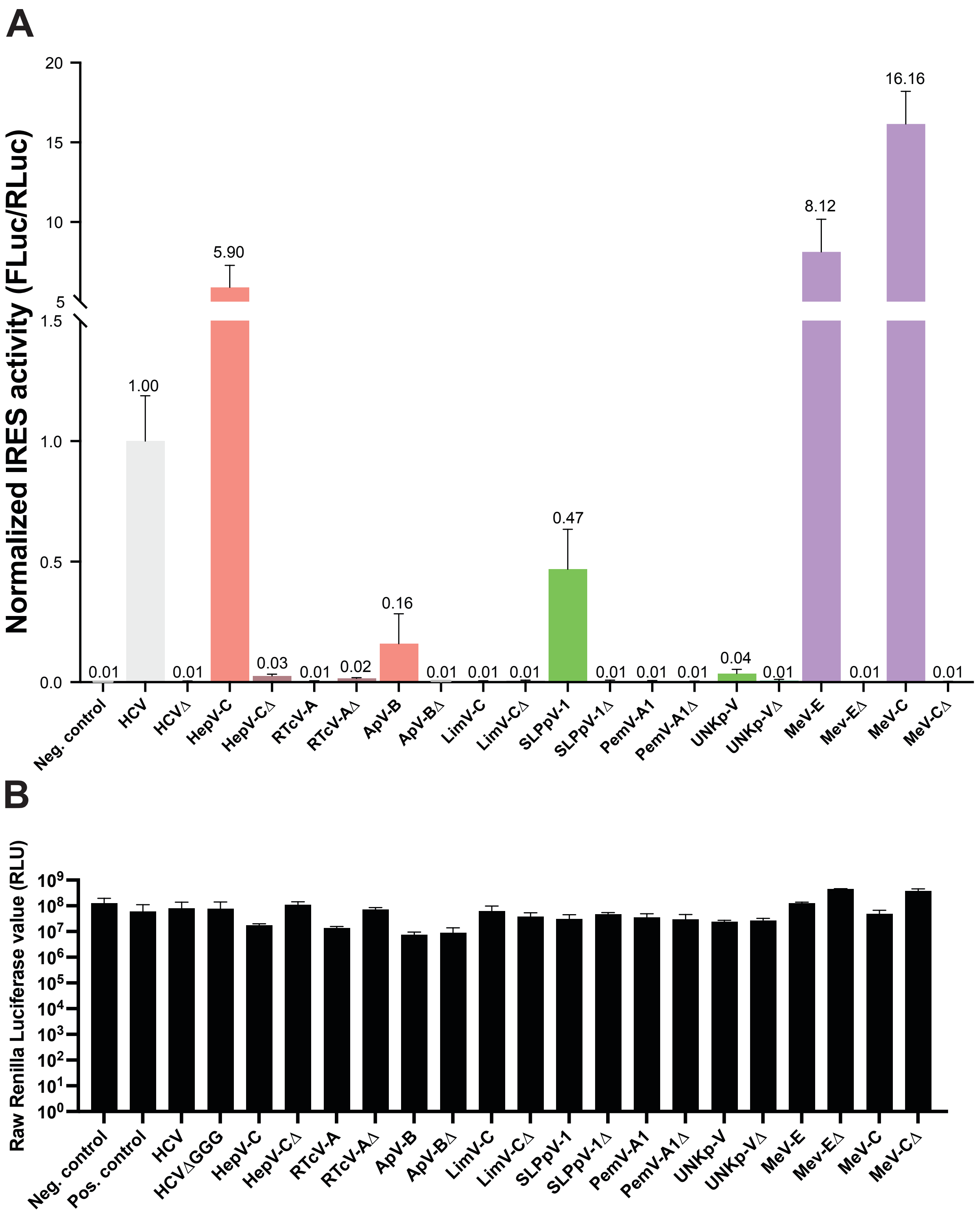


**Supplemental Figure S7.** Functional data on putative IRESs in a human *in vitro* translation system. In vitro translation assays of IRES activity using dual luciferase constructs in a homemade human lysate translation system (see methods). Dual luciferase constructs are as described in Figure 2. **(A)** The graph contains the results of translation assays of both wild-type and mutant (Δ) IRES sequences, normalized to wild type HCV IRES. The IRESs shown are as described in Figures 2 and S6. **(B)** Raw *Renilla* luciferase values for each construct tested in the human lysate translation system. Virus names and genomic information of IRESs are as described Figures 2 and S6.

| Accession number | Genomic Coordinates | Reason dropped |
|---|---|---|
| NC_021153 | 87-429 | ambiguious SS |
| MF352427 | 236-560 | ambiguious SS |
| NC_038314 | 237-576 | ambiguious SS |
| KY312542 | 37-275 | ambiguious SS |
| MH824542 | 39-356 | ambiguious SS |
| MT210612 | 145-541 | ambiguious seq |
| MT210615 | 6-342 | ambiguious seq |
| MT210611 | 95-491 | ambiguious seq |
| MT210606 | 11-335 | ambiguious seq |
| MT210613 | 71-397 | ambiguious seq |
| MT210607 | 168-491 | ambiguious seq |
| NC_012812 | 58-410 | ambiguious seq |

**Table S1. Sequences discarded from the covariation and clustering analysis.**
Of the 176 putative IRESs identified, 13 were dropped for further analysis due to ambiguous secondary structure (ambiguous SS) or degenerate sequencing (ambiguous seq).

| | *Flaviviridae* | *Picornaviridae* | *Caliciviridae* | *Picobirnaviridae* | *Unassigned* | ***Total*** |
|---|---|---|---|---|---|---|
| Cluster 1 | 56 | 35 | 1 | 1 | - | 93 |
| Cluster 2 | 3 | 33 | - | - | 12 | 48 |
| Cluster 3 | - | 21 | - | - | 1 | 22 |

| | *Mammalia* | *Aves* | *Reptilia* | *Aamphibia* | *Actinopteri* | *Insect* | *Unknown* | ***Total*** |
|---|---|---|---|---|---|---|---|---|
| Cluster 1 | 81 | 9 | - | - | - | 1 | 2 | 93 |
| Cluster 2 | 23 | 14 | 5 | 1 | 5 | - | - | 48 |
| Cluster 3 | - | 21 | 1 | - | - | - | - | 22 |

**Table S2. Taxonomic information on viral family and host class between Clusters.**

**Table S3. DNA oligonucleotide sequences used in this study.**

| DNA sequence | Description |
|---|---|
| GGGCCCTACGGGCCCTACTCGAGCTAAAGCTTGGC AATCCGGTACTGTTGGTAAAGCCACCGCCAGCCCC CTGATGGGGGCGACACTCCACCATGAATCACTCCC CTGTGAGGAACTACTGTCTTCACGCAGAAAGCGTCT AGCCATGGCGTTAGTATGAGTGTCGTGCAGCCTCCA GGACCCCCCCTCCCGGGAGAGCCATAGTGGTCTGC GGAACCGGTGAGTACACCGGAATTGCCAGGACGAC CGGGTCCTTTCTTGGATAAACCCGCTCAATGCCTGG AGATTTGGGCGTGCCCCCGCAAGACTGCTAGCCGA GTAGTGTTGGGTCGCGAAAGGCCTTGTGGTACTGC CTGATAGGGTGCTTGCGAGTGCCCCGGGAGGTCTC GTAGACCGTGCACCATGAGCACGAATCCTAAACCTC AAAGAAAAACCAAACGTAACACCAGATCTGAGGCAC GGCATAAGC | Gene block (dsDNA) fragment including the WT HCV IRES sequence with flanking restriction sites for cloning into pSGDLuc v3.0 to create dual luciferase reporter mRNAs |
| GGGCCCTACTCGAGCTAAAGCTTGGCAATCCGGTAC TGTTGGTAAAGCCACCGGACTAATTCCTATGTTTATA CGTTACTACCTTGTTCTGAACGGTGGGCGCCACCCC GCCTAGTAGGATCCTGGCTTATCGTGTAGACCTCTA GGGACCACATTAGCTAGAGTGTAGGCTGCTATGGAT GGAGTAGTGACCCCTTTTTGGGTATCACTCTCTAAG ACTCCGGAATGTGTCATAGTACGCTGGAAATCCTTAC TTGTTTTTCCATGAGGGGGAGGTGGTGCTGAAATAT TGCAAGCCACCCCTCGGTTAAAACAGTTTGGTGCC GCTTATGCCATATTACCGCCCCTTGTAGTTGGGCTGT TTTTGCAGCTCCGGGTTAGTAGAGTACCATAGTGGA CGCGGTGTTGGGAATCACCGCCTTGGCTGCACACT GCTTGATAGAGCTGCGGCTGGTCAAGCTAATTGTGG TATAACCAGTTGATTTGGCATATGGATTCTAGACTTAC CTTCTTACAAGATTTTCTTAAGGAACATAGATCTGAG GCACGGCATAAGC | Gene block (dsDNA) fragment including the WT MeV-E IRES sequence with flanking restriction sites for cloning into pSGDLuc v3.0 to create dual luciferase reporter mRNAs |
| GGGCCCTACGGGCCCTACTCGAGCTAAAGCTTGGC AATCCGGTACTGTTGGTAAAGCCACCTCGTATGCTAT CTCCTTGAACAATTTTTAATCATTGCGAAGTGATCATT GAAAAGGATAGGTGTTTAAGAACTCAAAGAGTGTTA ATAATGTTGGGTGACAGGTGTCCCCATAGAATTTATT AACATGATTTGGACTGGTTATCTAGTAAGAAGAACCA TCGAACGCACGAGCGAGCATTGCTTGCGGGGCAGT TACCCTGCGTCGATGTAAGTGTGTACCGGGGGGTG CACATGTTGATTCTTTATGGCCTGATAGGGTGCGTCA TTCGCGCCTAGATAATTAGTATAATGCGAATGGAATAA ATTTACATGGCTTCTATAATTGAAAATTTGACCACAAC TTTTGCATCATCAATGCTGGGAACAGCTGAGGATGC TGTCAGAAGATCTGAGGCACGGCATAAGC | Gene block (dsDNA) fragment including the WT LimV-C IRES sequence with flanking restriction sites for cloning into pSGDLuc v3.0 to create dual luciferase reporter mRNAs |
| GGGCCCTACGGGCCCTACTCGAGCTAAAGCTTGGC AATCCGGTACTGTTGGTAAAGCCACCTCCTACAAAT GCACATGAAGAACAGTTTGTAGAGATTAACAAACGC TTAGCTGAACCTAGGTGGTGAATCTAGTAGTAAGATA | Gene block (dsDNA) fragment including the HepV-C IRES sequence with flanking restriction sites for cloning into pSGDLuc |

| | |
|---|---|
| AGTAGAGGAAGCTATACCTTAAGTTGGTTGGGCCCT<br>CGTGTTTGCTCTATAAACAAAACCAAGTGAGTAGAGT<br>GGATGAACAGTACTAAATCCCTGAGTACAGGGAACC<br>TCACAGGTGTGATACACTTATGTCTATGTGACCTGGT<br>TGGAGGTTGGGCGTGCCCTATGATACTGGAGTGGG<br>AGATCTTTTGGGGAACCCACGTTTTCACACTGCCTG<br>ATAGGGTCTTGCCGAGAGACTCACTTGTTTCGGCTG<br>TACTTGTAACATGGAGAATAAAAATAAAGGAATTTTTC<br>AAACTGTTGGAGAGAGTTTGGATGGAATTTTGACTTT<br>GGCTGATAGAAGATCTGAGGCACGGCATAAGC | v3.0 to create dual luciferase reporter mRNAs |
| GGGCCCTACGGGCCCTACTCGAGCTAAAGCTTGGC<br>AATCCGGTACTGTTGGTAAAGCCACCTCGTATGCTAT<br>CTCCTTGAACAATTTTTAATCATTGCGAAGTGATCATT<br>GAAAAGGATAGGTGTTTAAGAACTCAAAGAGTGTTA<br>ATAATGTTGGGTGACAGGTGTCCCCATAGAATTTATT<br>AACATGATTTGGACTGGTTATCTAGTAAGAAGAACCA<br>TCGAACGCACGAGCGAGCATTGCTTGCGGGGCAGT<br>TACCCTGCGTCGATGTAAGTGTGTACCGCCCCGTGC<br>ACATGTTGATTCTTTATGGCCTGATAGGGTGCGTCAT<br>TCGCGCCTAGATAATTAGTATAATGCGAATGGAATAAA<br>TTTACATGGCTTCTATAATTGAAAATTTGACCACAACT<br>TTTGCATCATCAATGCTGGGAACAGCTGAGGATGCT<br>GTCAGAAGATCTGAGGCACGGCATAAGC | Gene block (dsDNA) fragment including the LimV-C ΔGGG IRES sequence with flanking restriction sites for cloning into pSGDLuc v3.0 to create dual luciferase reporter mRNAs |
| GGGCCCTACGGGCCCTACTCGAGCTAAAGCTTGGC<br>AATCCGGTACTGTTGGTAAAGCCACCTCCTACAAAT<br>GCACATGAAGAACAGTTTGTAGAGATTAACAAACGC<br>TTAGCTGAACCTAGGTGGTGAATCTAGTAGTAAGATA<br>AGTAGAGGAAGCTATACCTTAAGTTGGTTGGGCCCT<br>CGTGTTTGCTCTATAAACAAAACCAAGTGAGTAGAGT<br>GGATGAACAGTACTAAATCCCTGAGTACAGGGAACC<br>TCACAGGTGTGATACACTTATGTCTATGTGACCTGGT<br>TGGAGGTTGGGCGTGCCCTATGATACTGGAGTGGG<br>AGATCTTTTCCCGAACCCACGTTTTCACACTGCCTG<br>ATAGGGTCTTGCCGAGAGACTCACTTGTTTCGGCTG<br>TACTTGTAACATGGAGAATAAAAATAAAGGAATTTTTC<br>AAACTGTTGGAGAGAGTTTGGATGGAATTTTGACTTT<br>GGCTGATAGAAGATCTGAGGCACGGCATAAGC | Gene block (dsDNA) fragment including the HepV-C ΔGGG IRES sequence with flanking restriction sites for cloning into pSGDLuc v3.0 to create dual luciferase reporter mRNAs |
| GCTAGCCGAGTAGTGTTCCCTCGCGAAAGGCCTTG<br>TG | Forward ssDNA primer to introduce the ΔGGG into the HCV IRES sequence within pSGDLuc v3.0 via Q5 site-directed mutagenesis |
| AGTCTTGCGGGGGCACGC | Forward ssDNA primer to introduce the ΔGGG into the HCV IRES sequence within pSGDLuc v3.0 via Q5 site-directed mutagenesis |
| TAATACGACTCACTATAGGACTAATTCCTATGTTTATA<br>CGTTACTACCTTGTTCTG | Forward ssDNA primer to introduce the ΔGGG into the MeV-E IRES sequence within pSGDLuc v3.0 via Q5 site-directed mutagenesis |

| | |
|---|---|
| GTAAGAAGGTAAGTCTAGAATCCATATGCCAAATC | Reverse ssDNA primer to introduce the ΔGGG into the MeV-E IRES sequence within pSGDLuc v3.0 via Q5 site-directed mutagenesis |
| TAATACGACTCACTATAGGACTAATTCCTATGTTTATACGTTACTACCTTGTTCTG | Forward ssDNA primer to amplify MegE-V IRES templates for transcription of RNAs for SHAPE-MaP |
| GTAAGAAGGTAAGTCTAGAATCCATATGCCAAATC | Reverse ssDNA primer to amplify MegE-V IRES templates for transcription of RNAs for SHAPE-MaP |
| TAATACGACTCACTATAGGCTCCTACAAATGCACATGAAGAACAGTTTGTAG | Forward ssDNA primer to amplify HepV-C IRES templates for transcription of RNAs for SHAPE-MaP |
| AAGTCAAAATTCCATCCAAACTCTCTCCAAC | Reverse ssDNA primer to amplify HepV-C IRES templates for transcription of RNAs for SHAPE-MaP |
| TAATACGACTCACTATAGGGTTTCTGAGCACTGGTAAGAGCTTAGAC | Forward ssDNA primer to amplify LimV-C IRES templates for transcription of RNAs for SHAPE-MaP |
| TGATGATGCAAAAGTTGTGGTCAAATTTTCAATTATAG | Reverse ssDNA primer to amplify LimV-C IRES templates for transcription of RNAs for SHAPE-MaP |
